# Oligodendrocyte precursor cells exacerbate acute CNS inflammation via macrophage and T cell activation in a mouse model of multiple sclerosis

**DOI:** 10.1101/2024.05.28.596190

**Authors:** Kana Ohashi, Nagi Uemura, Kazuki Nagayasu, Shuji Kaneko, Takakuni Maki, Hisashi Shirakawa

## Abstract

Oligodendrocyte precursor cells (OPCs) are a type of glial cell that differentiates into mature oligodendrocytes, a cell type that contributes to myelination, but their roles in the pathologies are not fully understood. Activities other than differentiation into oligodendrocytes have recently been reported for OPCs present in the inflammatory milieu, but intervention studies using animal models are lacking. This study aimed to explore the role of OPCs in mouse MS model experimental autoimmune encephalomyelitis (EAE). Using inducible diphtheria toxin receptor-expressing transgenic mice, platelet-derived growth factor receptor A (PDGFRα)^+^ OPCs were depleted in EAE mice. Surprisingly, OPC depletion in the acute phase improved clinical scores and reduced demyelination. Major histocompatibility complex (MHC) class II was reduced in the spinal cord, whereas astrocyte marker and blood–spinal cord barrier tight junction and adhesion molecule expressions were unaffected after OPC depletion. The numbers of T cells and IL17-expressing Th17 cells were decreased in the spinal cords of the OPC-depleted group. MHC class II expression in spinal cord macrophages was consistently decreased by OPC depletion. These data suggest that in the acute phase of EAE, OPCs are involved in activation of infiltrated macrophages and induce subsequent T cell activation and neuroinflammation. Although the precise mechanisms remain unclear, this implies that OPCs exist not only as the source for oligodendrocytes but also play a pivotal role in central nervous system (CNS) autoimmune inflammation.

**Table of Contents Image:** 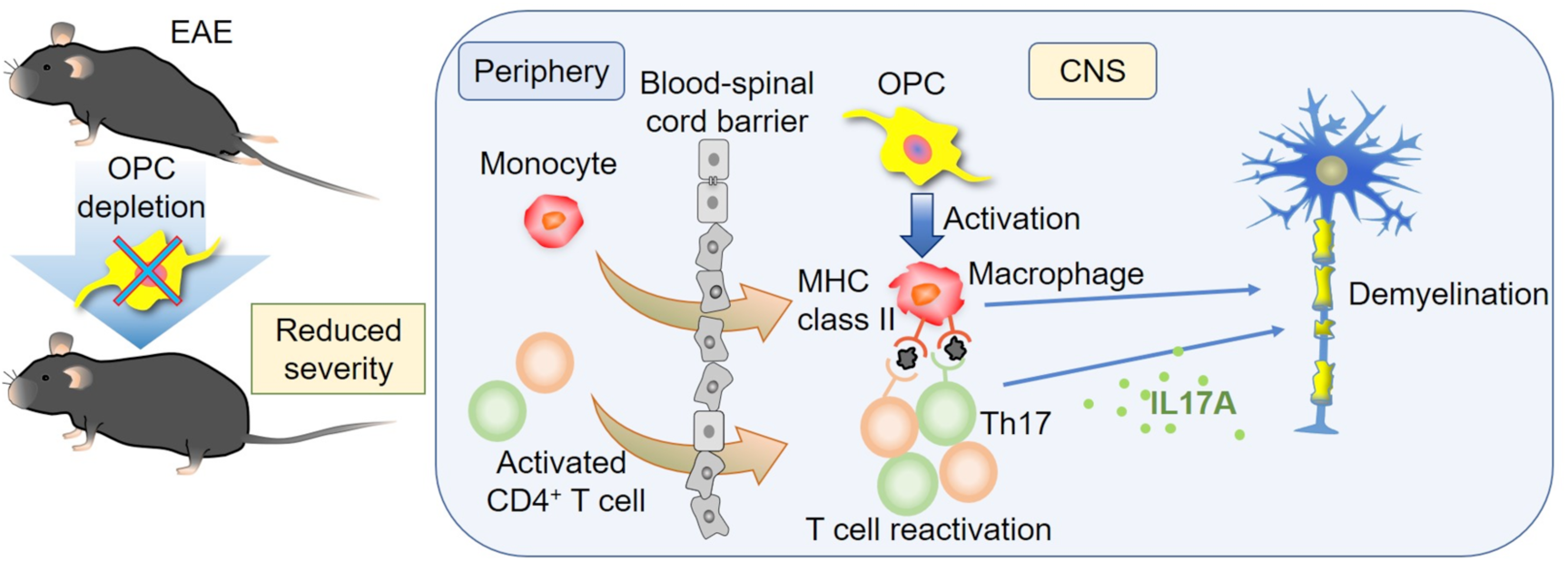

**MAIN POINTS:**

- OPC depletion in the acute phase of EAE improved clinical scores and reduced demyelination.
- OPC depletion in the spinal cord suppressed Antigen presentation via MHC class II.
- OPCs are involved in activation of infiltrated macrophages and induce subsequent T cell activation and neuroinflammation.

## 1 INTRODUCTION

Multiple sclerosis (MS) is an inflammatory demyelinating disease of the central nervous system (CNS) where autoreactive T cells and macrophages infiltrate the CNS and provoke widespread inflammation and subsequent demyelination. In the CNS, antigen-specific CD4^+^ T cells are reactivated by antigen-presenting cells (Chastain et al., 2011; Dong and Yong, 2019), proliferate by clonal expansion (Planas et al., 2018) and produce inflammatory cytokines such as IL17. However, the specific interactions that occur between these peripheral immune cells and resident CNS cells remain unclear.

Oligodendrocyte precursor cells (OPCs) are a type of glial cell that differentiates into mature oligodendrocytes. Oligodendrocytes produce myelin sheaths and provide trophic supports to axons (Fünfschilling et al., 2012) in a healthy CNS. As OPCs are a source of myelin producing cells, they are presumed to be a protective cell type that contributes to myelin repair in MS. However, differentiation capacity of OPCs is limited in the MS lesions (Franklin and Ffrench-Constant, 2017; Nicaise et al., 2019). ^14^C analysis of brains from patients with MS suggests that in the majority of individuals, remyelination seems to be conducted mainly by pre-existing oligodendrocytes and not by new oligodendrocytes differentiated from OPCs (Yeung et al., 2019). Furthermore, OPCs have been shown to be epigenetically suppressed in their differentiation under MS pathology (Tiane et al., 2023; Liu et al., 2024). These raise the possibility that most OPCs have functions beyond the replacement of oligodendrocytes.

Several transcriptomic analyses have revealed that OPCs in patients with MS and experimental autoimmune encephalomyelitis (EAE) mice change their phenotypes to upregulate immune-related molecules (Falcão et al., 2018; Jäkel et al., 2019; Kirby et al., 2019; Meijer M, et al., 2022). However, these experiments are gene expression analyses or in vitro studies, and the lack of in vivo studies leaves the contribution of immature OPCs to EAE pathology largely unclear, especially in the acute phase, when infiltration of peripheral immune cells and demyelination begin.

In the present study, OPCs were depleted during EAE to examine their role in this disease. We used transgenic mice in which expression of the diphtheria toxin receptor (DTR) can be induced under the control of the *Pdgfra* promoter. Platelet-derived growth factor receptor A (PDGFRα) is a receptor tyrosine kinase involved in OPC survival and proliferation (Barres et al., 1992). In this system, upon administration of diphtheria toxin, DTR-expressing cells are depleted. We found that depletion of OPCs during the acute phase of EAE improves clinical symptoms. Therefore, we investigated the effect of OPC depletion on the cellular dynamics in EAE pathology and found that the number of T cells infiltrating/proliferating was reduced in the CNS. We then analyzed this phenomenon in more detail.

## 2 MATERIALS AND METHODS

### 2.1 Animals

B6N.Cg-Tg(Pdgfra-cre/ERT)467Dbe/J (Jackson strain #018280) were crossed with C57BL/6-Gt(ROSA)26Sor^tm1(HBEGF)Awai^/J (Jackson strain #007900) mice to generate *Pdgfra^CreER/+^;Rosa26^DTR/+^*and *Pdgfra^+/+^;Rosa26^DTR/+^* mice. For anesthesia, a mixed solution of medetomidine, midazolam, and butorphanol was used. All experiments were conducted in accordance with the ethical guidelines of the Kyoto University Animal Research Committee.

### 2.2 PDGFR*α*^+^ cell depletion

A total of 150 mg/kg/day Tamoxifen (T5648, Sigma-Aldrich) in corn oil (C8267, Sigma-Aldrich) and 2–3 µg/kg/day diphtheria toxin (D0564, Sigma-Aldrich) were intraperitoneally injected into *Pdgfra^CreER/+^;Rosa26^DTR/+^*(Depl) and *Pdgfra^+/+^;Rosa26^DTR/+^* (Ctrl) mice. For OPC depletion in the I mice (Figure. 2), male and female 2–3-month-old mice were used.

**Figure 1.**
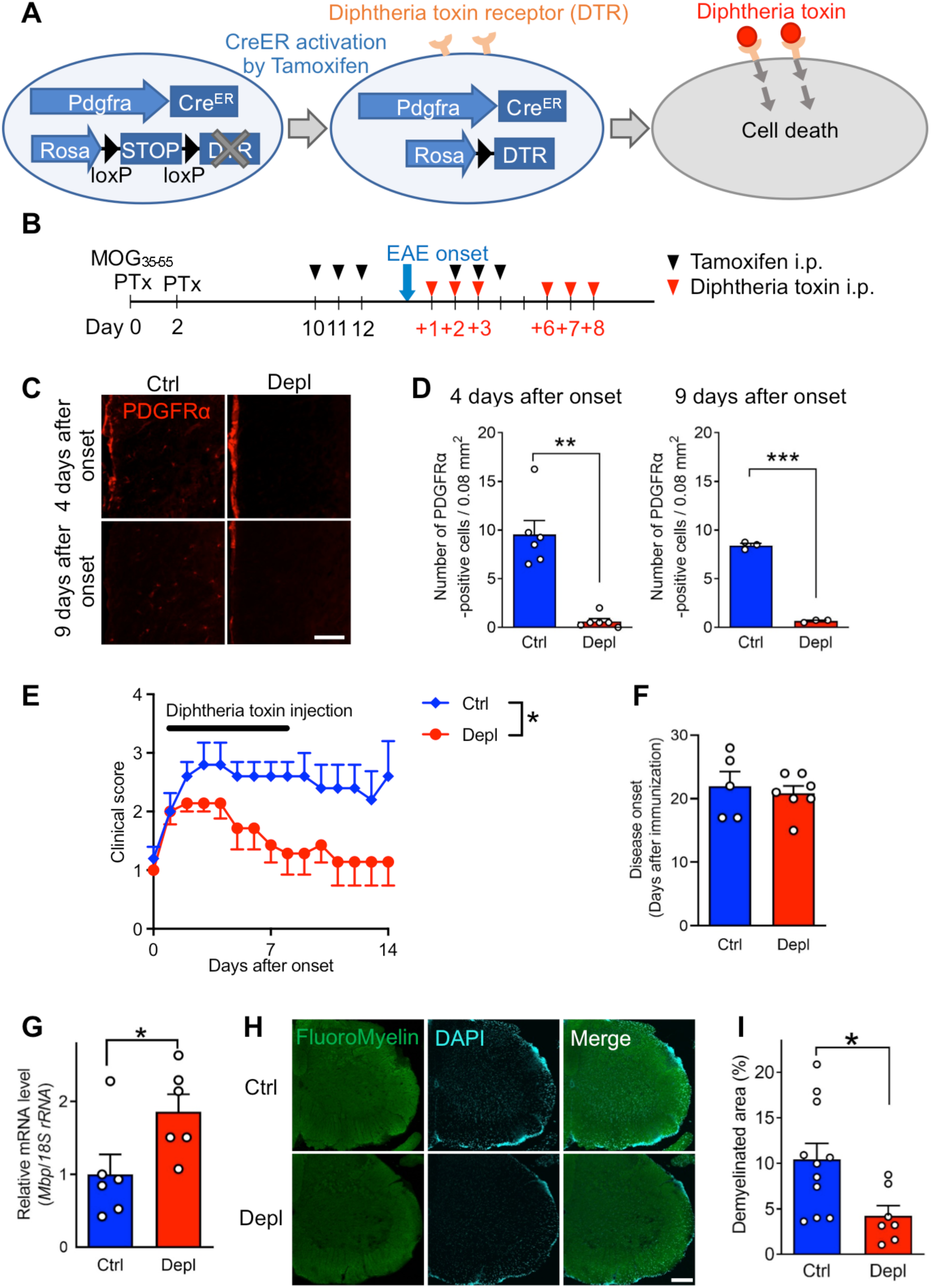
EAE clinical score and demyelination are reduced after OPC depletion in the acute phase. ***A***. Schematic diagram of CreER-mediated expression of diphtheria toxin receptor (DTR) and depletion of PDGFRα^+^ cells after diphtheria toxin administration in *Pdgfra^CreER/+^;Rosa26^DTR/+^*mice. ***B-I***. Depletion of PDGFRα^+^ cells in the acute phase of EAE. ***B***. Time course of EAE induction and intraperitoneal injection of tamoxifen and diphtheria toxin. ***C, D***. Representative images of PDGFRα immunostaining in the lumbar spinal cord at 4 and 9 days after onset (***C***), and quantification of number of expressing cells in 0.08 mm^2^ (***D***). Scale bar, 100 µm. *n* = 6 mice for 4 days after onset; *n* = 3 mice for 9 days after onset. ***E***. EAE clinical score after disease onset. *n* = 5–7 mice. Two-way ANOVA with Bonferroni’s multiple comparison test. ***F***. Days before symptom onset after immunization. *n* = 5–7 mice. ***G***. Quantitative PCR analysis of *Mbp* in spinal cords collected at 4 days after EAE onset (after three injections of diphtheria toxin). *n* = 6 mice. ***H, I***. Representative images of FluoroMyelin (myelinated area) and DAPI (cell nuclei) staining (***H***), and quantification of percentage of demyelinated area (***I***) in the spinal cord white matter at 4 days after the onset of EAE. Scale bar, 200 µm. *n* = 7–11 mice. \**P* < 0.05, \*\**P* < 0.01, \*\*\**P* < 0.001. Data are expressed as mean ± SEM.

**Figure 2.**
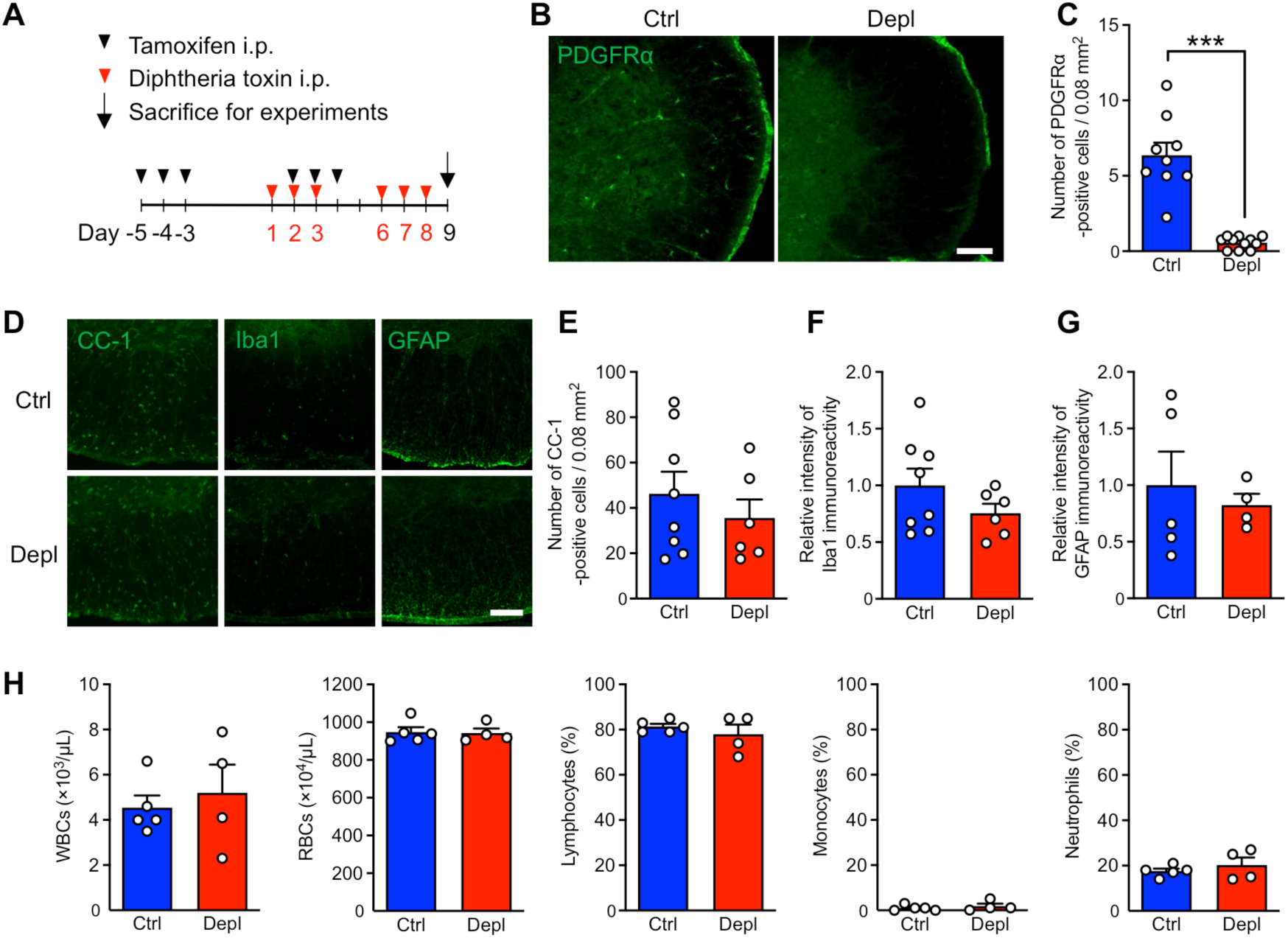
OPC depletion in naïve mice does not affect glial cell and blood cell composition. ***A***. Time course of OPC depletion in naïve mice. ***B, C***. Fluorescent immunostaining for PDGFRα in the lumbar spinal cord at day 9 (***B***) and quantification of the number of positive cells in 0.08 mm^2^ (***C***). Scale bar, 100 µm. *n* = 9–11 mice. ***D-G***. Fluorescent immunostaining for CC-1, Iba1 and GFAP in the lumbar spinal cord at day 9. Scale bar, 100 µm. Quantification of the number of CC-1 positive cells (***E***) in 0.08 mm^2^.area, relative fluorescence intensity of Iba1 (***F***) and GFAP (***G***) in the white matter area. *n* = 6–8 mice. ***H***. The numbers of white blood cells (WBCs), red blood cells (RBCs), and the percentages of lymphocytes, monocytes, and neutrophils in the WBCs at day 9. *n* = 4–5 mice. \*\*\**P* < 0.001. Data are expressed as mean ± SEM.

### 2.3 EAE

Active EAE was induced in female 7–12-week-old mice as described previously (Miyamura et al., 2019), with minor modifications. Briefly, *Pdgfra^CreER/+^;Rosa26^DTR/+^*(Depl) and *Pdgfra^+/+^;Rosa26^DTR/+^* (Ctrl) mice were immunized subcutaneously with 200 µl of the emulsion, a 1:1 mixture of 200 µg of myelin oligodendrocyte glycoprotein peptide 35–55 (MOG 35-55; Scrum) in saline and complete Freund’s adjuvant (231131, BD Difco) containing 6 mg/ml *Mycobacterium tuberculosis* H37Ra (231141, BD Difco). Mice received 17.5 µg/kg pertussis toxin (P7208, Sigma-Aldrich) intraperitoneally on the day of immunization and 2 days later. For the immunization of wild-type (WT) mice, an emulsion containing 100 µg of MOG 35-55 and 10 µg/kg pertussis toxin was injected. The clinical score for the mice was measured every other day until 10 days post immunization and every day thereafter. Scoring was as follows: 0, no clinical deficit; 1, partial tail paralysis; 2, full tail paralysis; 3, partial hindlimb paralysis; 4, full hindlimb paralysis; and 5, forelimb paresis. Only mice that developed EAE symptoms 14–28 days after immunization were analyzed. Mice that developed atypical EAE symptoms (strong head tilt) were excluded.

### 2.4 Blood test

The mice were anesthetized and decapitated, and whole blood was collected in tubes containing EDTA-2K. The blood samples were sent to the Sanritsu Zelkova Veterinary Laboratory (Tokyo) via refrigerated shipping.

### 2.5 Quantitative RT-PCR

Mice were anesthetized and transcardially perfused with phosphate buffered saline (PBS). The lumbar spinal cord (L3–L5), subiliac lymph node, and spleen were harvested 4 days post EAE onset. Total RNA was extracted using an ISOGEN kit (Wako). First-strand cDNA was prepared using a ReverTra Ace qPCR RT Kit (Toyobo). Quantitative PCR was performed using the StepOne Real-Time PCR System (Applied Biosystems) and the THUNDERBIRD SYBR qPCR Mix (Toyobo). The primer sets used were as follows: 5′-GCA ATT ATT CCC CAT GAA CG-3′ and 5′-GGC CTC ACT AAA CCA TCC AA-3′ for 18s ribosomal RNA (18s rRNA); 5′-TGT ACA AGG ACT CAC ACA CGA-3′ and 5′-CTT CCC TTG GGA TGG AGG TG-3′ for *Mbp*; 5′-AGT TAA GGC ACG GGT AGC AC-3′ and 5′-CAA CGA TGT TGG CGA ACC AG-3′ for *Cldn5*; 5′-GCT GCC TCG AAC CTC TAC TC-3′ and 5′-ACT GCT TGG GCT CAG ATG AC -3′ for *Tjp1*; 5′-TAC TGT TTG CAG TCT CTC AAG C-3′ and 5′-TCA CCT TCG CGT TTA GTG GG-3′ for *Vcam1*; 5′-AGC TCG GAG GAT CAC AAA CG-3′ and 5′-TCC AGC CGA GGA CCA TAC AG-3′ for *Icam1*; 5′-GTA AAG CGT GAA GAC AGC TGC-3′ and 5′-CTG AAC CCA AGC TCA CAG G-3′ for *H2-D1*; 5′-GTG CGG GGG ACC CAA AGA CCA AAC-3′ and 5′-GCA CGT GGA GGT GAA CCA TCC TTA TAT-3′ for *Gzmb*; 5′-AGC CTC TGT GGA GGT G-3′ and 5′-TAC TGG CCA ATG TCT C-3′ for *H2-Aa*; 5′-CTG GAT GAA GCA GTG GCT CT-3′ and 5′-CCC AGG CCA GAA GAT AGG TC-3′ for *Cd74*; 5′-GGA TAA GGT CGA AGA GGA AGT AA-3′ and 5′-CGT TAT ATC CTT GAC AAG GTC TT-3′ for *Cd40l*g; 5′-CGG AGC GGA CCA ACA GCA TCG TTT C-3′ and 5′-CAG GGT AGC CAT CCA CGG GCG GGT-3′ for *Tbet*; 5′-GGC AGA ACC GGC CCC TTA TC-3′ and 5′-TGG TCT GAC AGT TCG CGC AG-3′ for *Gata3*; 5′- GAC AGG GAG CCA AGT TCT CA-3′ and 5′- CTT GTC CCC ACA GAT CTT GCA-3′ for *Rorc*; 5′-CCT GGT TGT GAG AAG GTC TTC G-3′ and 5′-TGC TCC AGA GAC TGC ACC ACT T-3′ for *Foxp3*; The PCR conditions were as follows: 95 °C for 10 min, followed by 40 cycles of 95 °C for 15 s and 60 °C for 60 s. Expression of each gene was normalized against that of 18S rRNA.

### 2.6 Immunohistochemistry

Mice were anesthetized and transcardially perfused with PBS and 4 % paraformaldehyde (PFA). The extracted spinal cord (L3–L5) was postfixed for 3 h in 4 % PFA, placed in 15 % sucrose in 0.1 M phosphate buffer, and then embedded in OCT compound. Coronal sections (20 µm thick) were cut using a cryomicrotome (Leica), blocked with PBS containing 3 % bovine serum albumin, and permeabilized with 0.1 % Triton X-100 in blocking solution. Sections were incubated with primary antibodies at 4 °C overnight (14-20 h) at the following dilutions: anti-PDGFRα (AF1062-SP, goat IgG, 1:250; R&D Systems), anti-APC (Ab-7) Mouse mAb (CC-1) (OP80, mouse IgG, 1:200; Calbiochem®), anti-ionized calcium binding adaptor molecule 1 (Iba1) (019-19741, rabbit IgG, 1:200; Wako Pure Chemical Industries), anti-glial fibrillary acidic protein (GFAP) (ab7260, rabbit IgG, 1:200; Abcam), anti-CD31 (553370, rat IgG, 1:200; BD Biosciences), anti-CD3 (555273, rat IgG, 1:200; BD Biosciences), anti-IL17 (NBP1-76337, rabbit IgG, 1:150; Novus Biologicals), anti-MHC class II (556999, rat IgG, 1:50; BD Biosciences), anti-collagen I (ab21286, rabbit IgG, 1:250; Abcam), anti-CD45 (AF114-SP, goat IgG, 1:50; R&D Systems), anti-CD11c (14-0114-82, Armenian hamster IgG, 1:50; eBioscience). After washing, the sections were incubated with the following fluorescence-labeled secondary antibodies at room temperature (22-28 °C) for 1.5 h at different dilutions: Alexa Fluor 488-labeled donkey anti-goat IgG (1:300; A11055) or anti-mouse IgG (1:300; A21202) or anti-rabbit IgG (1:500; A21206) or anti-rat IgG (1:500; A21208, both Invitrogen), Alexa Fluor 594-labeled donkey anti-goat IgG (1:300; A11058) or anti-rabbit IgG (1:500; A21207, both Invitrogen), DyLight 594-labeled goat anti-Armenian hamster IgG (1:200; 405512, BioLegend), and Alexa Fluor 647-labeled donkey anti-rabbit IgG (1:500; A31573, Invitrogen). Images were acquired using FluoView FV10i confocal microscope (Olympus). The number of PDGFRα^+^ and CC-1^+^ cells in a 0.08 mm^2^ field of the white matter, the number of CD3^+^ and CD3^+^IL17A^+^ cells per image, and the intensities of the GFAP-, CD31-, Iba1-, and MHC class II-immunofluorescent signals in the white matter were quantified using ImageJ software.

### 2.7 FluoroMyelin™ staining

Demyelination was evaluated as previously described (Tsutsui et al., 2018), with minor modifications. Briefly, frozen coronal spinal cord sections (20 µm thick) were stained with FluoroMyelin green fluorescent myelin stain (1:1000, Invitrogen) and mounted with DAPI Fluoromount-G (Southern Biotech). The demyelinated and total areas of white matter were measured using ImageJ software. The percentage of demyelinated areas was calculated as follows: demyelinated area (%) = (demyelinated area in white matter) / (total white matter area) × 100 (%).

### 2.8 Experimental design and statistical analysis

Statistical analysis was performed using GraphPad Prism 9 software. Details of the procedures used for statistical analyses (including which tests were performed, exact p-values, and sample sizes) and details about experimental design are provided in the Results section or in the legend to each figure. For comparisons between a single experimental group and a control group, an unpaired Student’s *t*-test or Welch’s *t*-test was used. In addition, a two-way analysis of variance (ANOVA) with Bonferroni’s post hoc test was used, as appropriate. All data are expressed as the mean ± SEM. Statistical significance was set at *P* < 0.05.

## 3 RESULTS

### 3.1 Depletion of PDGFR*α*^+^ OPCs in acute phase of EAE ameliorated clinical score and demyelination

We used *Pdgfra^CreER/+^;Rosa26^DTR/+^* mice to deplete OPCs. In these mice, after tamoxifen injection, CreER activation and subsequent diphtheria toxin receptor expression was selectively induced in PDGFRα^+^ cells. Upon administration of diphtheria toxin, the PDGFRα^+^ cells were depleted (Fig. 1A). *Pdgfra^+/+^;Rosa26^DTR/+^* mice were used as the controls as these mice do not express CreER and administration of diphtheria toxin do not deplete PDGFRa^+^ cells. To minimize the depletion of mature oligodendrocytes differentiated from PDGFRα^+^ OPCs, tamoxifen injection was started 10 days post immunization. Mice were injected with diphtheria toxin the day after symptom onset (Fig. 1B). Efficient depletion of PDGFRα^+^ OPCs in the spinal cord was validated by immunostaining at 4 and 9 days after the onset of EAE (Fig. 1C, D; *t*_(10)_*=* 6.086, *P* = 0.0013, Unpaired Welch’s *t-*test for 4 days after the onset; *t*_(4)_*=* 32.88, *P* < 0.0001, Unpaired Student’s *t-*test for 9 days after the onset). The EAE clinical score was significantly suppressed in the OPC-depleted group (Fig. 1E; *F*_(1,10)_ = 6.604, *P* = 0.0279, Two-way ANOVA), whereas the day of onset did not vary between the groups (Fig. 1F; *t*_(10)_*=* 0.4897, *P* = 0.6349, Unpaired Student’s *t-*test). In accordance with the reduced severity of EAE after OPC depletion in the acute phase, the mRNA expression level of *Mbp,* the gene for myelin protein, was significantly upregulated in the OPC-depleted group (Fig. 1G; *t*_(10)_*=* 2.367, *P* = 0.0395, Unpaired Student’s *t-*test). The demyelinated area was significantly reduced in the OPC-depleted group (Fig. 1H, I; *t*_(16)_*=* 2.558, *P* = 0.0211, Unpaired Student’s *t-*test). After the depletion of OPCs in naïve mice (Fig. 2A-C; *t*_(18)_*=* 7.536, *P* = 0.0001, Welch’s *t-*test), glial cell (Fig. 2D-G; *t*_(12)_*=* 0.7994, *P* = 0.4396, Unpaired Student’s *t-*test for the oligodendrocyte marker CC-1; *t*_(12)_*=* 1.305, *P* = 0.2163, Unpaired Student’s *t-*test for the microglia marker Iba1; *t*_(7)_*=* 0.5081, *P* = 0.6270, Unpaired Student’s *t-*test for the astrocyte marker GFAP) and the peripheral blood cell remained unchanged (Fig. 2H; *t*_(7)_*=* 0.5258, *P* = 0.6153, Unpaired Student’s *t-*test for WBC; *t*_(7)_*=* 0.1456, *P* = 0.8883, Unpaired Student’s *t-*test for RBC; *t*_(7)_*=* 0.7761, *P* = 0.4873, Unpaired Welch’s *t-*test for lymphocytes; *t*_(7)_*=* 0.6494, *P* = 0.5368, Unpaired Student’s *t-*test for monocytes; *t*_(7)_*=* 0.8264, *P* = 0.4358, Unpaired Student’s *t-*test for neutrophils), indicating that these cells were not affected by this protocol. PDGFRα expression has also been reported in collagen-expressing fibroblasts (Dorrier et al., 2021), and in accordance with previous studies (Yahn et al., 2020; Dorrier et al., 2021), we observed collagen I^+^PDGFRα^+^ fibroblasts in chronic lesions (Fig. 3A). In acute lesions, a relatively small number of collagen I^+^ cells were present, but the PDGFRα^+^ cells were predominantly collagen I^−^ OPCs (Fig. 3B). These data indicate that OPC depletion during the acute phase of EAE suppresses the severity of EAE symptoms and demyelination.

**Figure 3.**
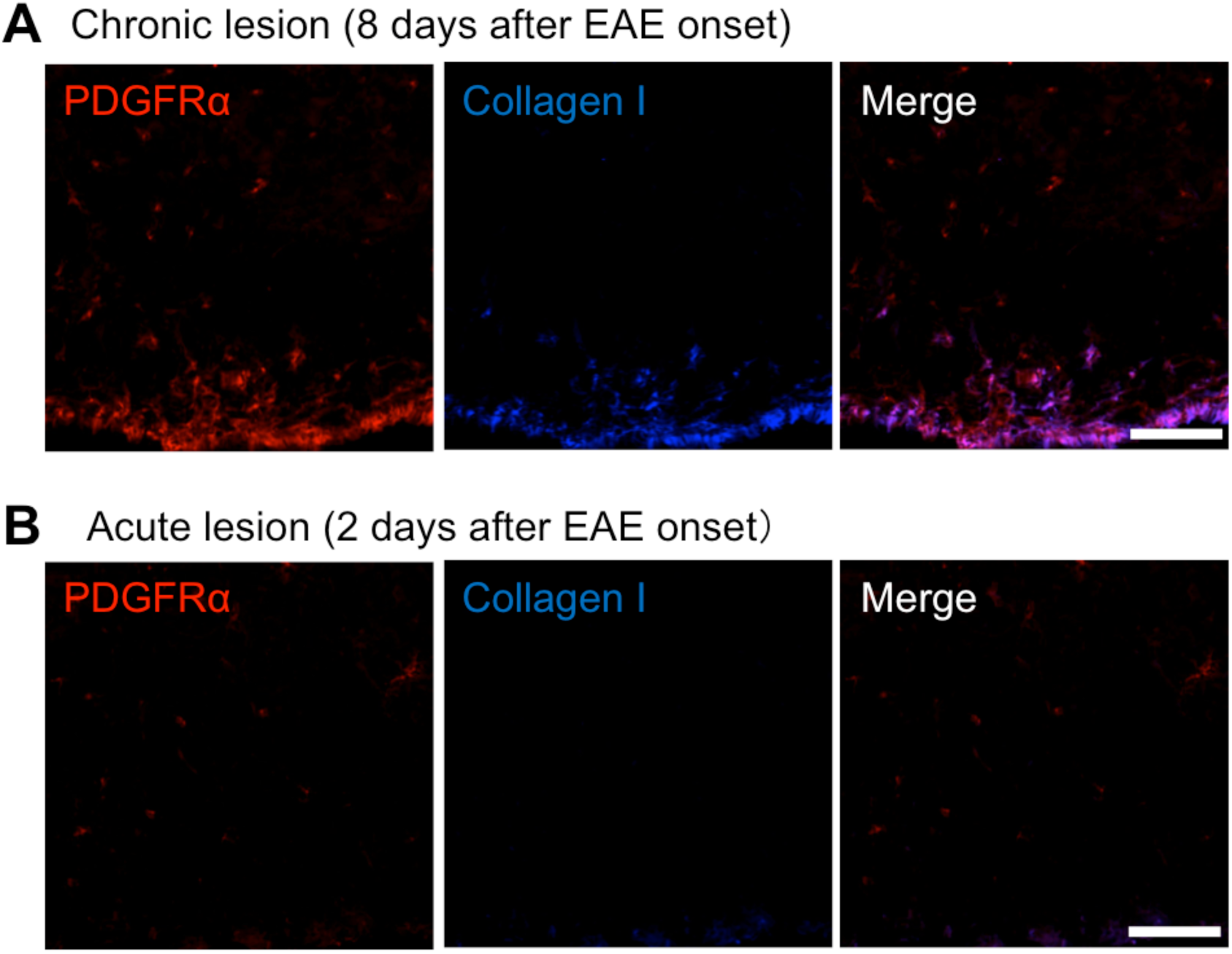
PDGFR*α* is mainly expressed in OPCs but also expressed in fibroblasts in chronic fibrotic scars. Co-immunostaining of PDGFRα, and collagen I during the chronic (8 days after onset, ***A***) and acute phases of EAE (2 days after onset, ***B***) in the spinal cord white matter lesions of wild-type mice. Scale bar, 100 µm.

### 3.2 Antigen presentation via MHC class II is suppressed in the spinal cord of OPC-depleted mice

To explore the mechanisms involved in the amelioration of EAE by OPC depletion, immunostaining and quantitative PCR analysis were conducted in spinal cord samples 4 days after EAE onset. Astrocytes are the most abundant cells in the CNS and contribute to inflammation through the production of pro-inflammatory cytokines. However, the fluorescent intensity of the astrocyte marker GFAP in the white matter did not change between the groups (Fig. 4A, B; *t*_(10)_*=* 1.202, *P* = 0.2572, Unpaired Student’s *t-*test). Peripheral immune cells infiltrate the CNS through the disrupted blood–brain barrier (BBB) and blood–spinal cord barrier (BSCB), and interact with adhesion molecules on endothelial cells. Previous studies indicate that a subset of OPCs is localized near the blood vessels and induces BBB/BSCB breakdown in MS and ischemic injury models (Seo et al., 2013; Maki, 2017; Niu et al., 2019). However, there were no significant differences in the expression levels of the endothelial cell marker protein CD31 (Fig. 4C, D; *t*_(4)_*=* 0.1807, *P* = 0.8654, Unpaired Student’s *t-*test), tight junction protein mRNAs *Cldn5* and *Tjp1* (Fig. 4E; *t*_(10)_*=* 1.628, *P* = 0.1346, Unpaired Student’s *t-*test; Fig. 4F; *t*_(10)_*=* 0.5913, *P* = 0.5675, Unpaired Student’s *t-*test), or adhesion molecule mRNAs *Vcam1* and *Icam1* (Fig. 4G; *t*_(10)_*=* 0.6461, *P* = 0.5328, Unpaired Student’s *t-*test; Fig. 4H; *t*_(10)_*=* 1.467, *P* = 0.1902, Unpaired Welch’s *t-*test). These results suggest that OPC depletion does not affect astrocyte activation and BSCB function.

**Figure 4.**
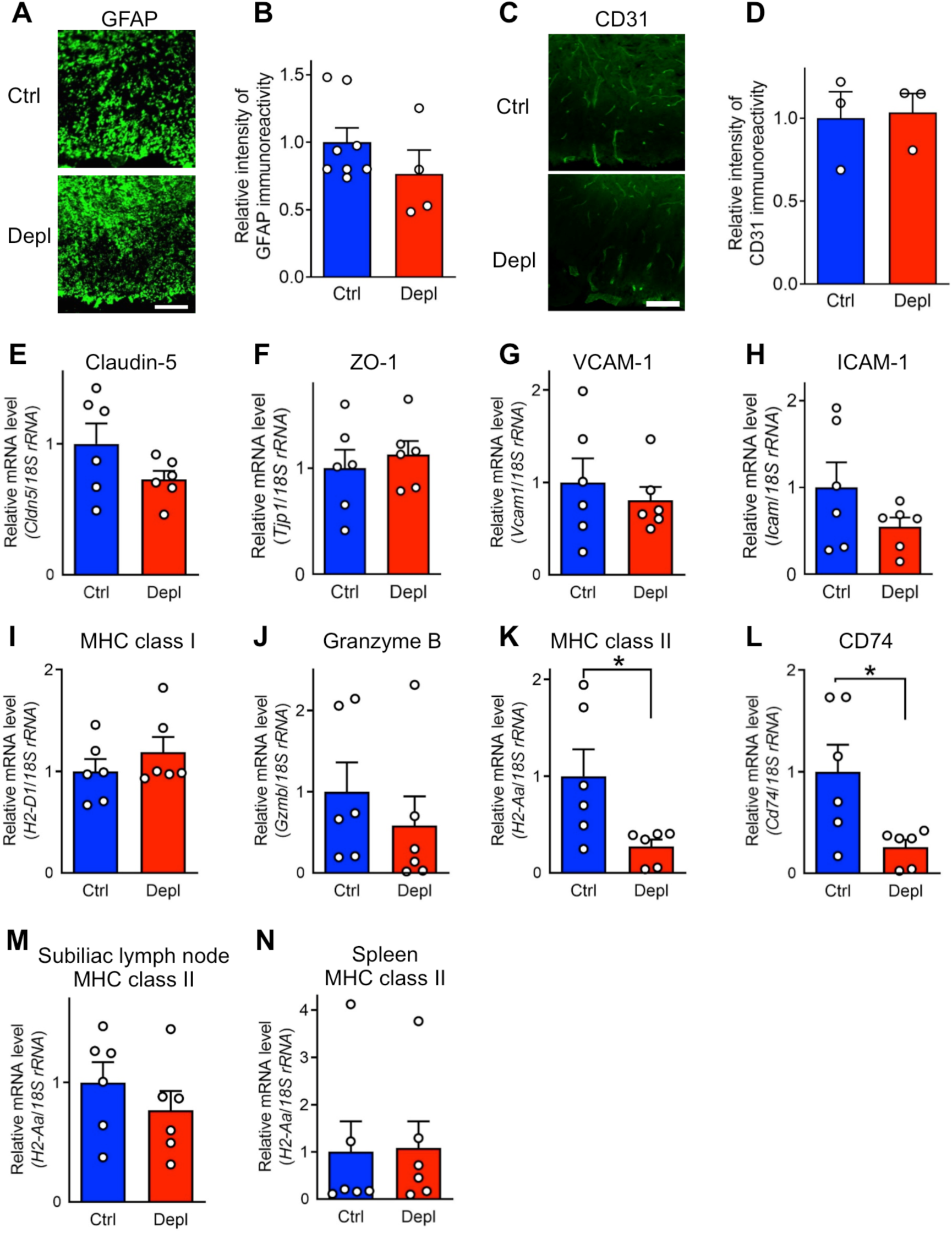
MHC class II is reduced in the spinal cord, whereas astrocyte marker and blood– spinal cord barrier tight junction and adhesion molecule expressions are unaffected after OPC depletion in the acute phase. *A, B*. Representative immunostaining images of GFAP (*A*) and quantification of relative fluorescence intensity in the white matter area at 4 days after onset of EAE (*B*). Scale bar, 100 µm. *n* = 4–8 mice. *C, D*. Representative immunostaining images of CD31 (*C*) and quantification of relative fluorescence intensity in the white matter area at 4 days after onset of EAE (*D*). Scale bar, 100 µm. *n* = 3 mice. *E-L*. Quantitative PCR analysis of *Cldn5* (*E*), *Tjp1* (*F*), *Vcam1* (*G*), *Icam1* (*H*), MHC-I *H2-D1* (*I*), *Gzmb* (*J*), MHC class II *H2-Aa* (*K*), and *Cd74* (*L*) in the spinal cord 4 days after onset. *n* = 6 mice. *M, N*. Quantitative PCR analysis of the MHC class II gene *H2-Aa* in the subiliac lymph node (*M*) and spleen (*N*) 4 days after onset. *n* = 6 mice. \**P* < 0.05. Data are expressed as mean ± SEM.

Pathogenic CD4^+^ and CD8^+^ T-cell responses play pivotal roles in EAE. Peripherally activated T cells require reactivation by antigen presentation in the CNS to mediate encephalitogenic activity (Kawakami et al., 2004). There were no differences in the expression levels of the MHC-I gene *H2-D1* (Fig. 4I; *t*_(10)_*=* 0.9968, *P* = 0.3424, Unpaired Student’s *t-*test) and the cytotoxic serine protease that is released by CD8^+^ T cells, Granzyme B (*Gzmb*) (Fig. 4J; *t*_(10)_*=* 0.8168, *P* = 0.4331, Unpaired Student’s *t-*test). In contrast, the expression levels of the MHC class II gene *H2-Aa* (Fig. 4K; *t*_(10)_*=* 2.526, *P* = 0.0472, Unpaired Welch’s *t-*test) and its chaperone *Cd74* (Fig. 4L; *t*_(10)_*=* 2.693, *P* = 0.0377, Unpaired Welch’s *t-*test) were significantly decreased in the spinal cord samples from the OPC-depleted group. *H2-Aa* expression was unaltered in the subiliac lymph nodes and spleen, indicating that antigen presentation was selectively reduced in the CNS (Fig. 4M; *t*_(10)_*=* 0.9906, *P* = 0.3452, Unpaired Student’s *t-*test; Fig. 4N; *t*_(10)_*=* 0.09560, *P* = 0.9257, Unpaired Student’s *t-*test). These data imply that in the spinal cord, antigen presentation via MHC class II is suppressed via OPC depletion.

3.3 **OPC depletion resulted in suppressed Th17 response and effector cytokine expression**

Consistent with the apparent reduced antigen presentation via MHC class II (Fig. 4K), *Cd40lg* (CD40L/CD154), which is highly expressed on activated CD4^+^ T cells, was significantly decreased in the spinal cord samples from the OPC-depleted group (Fig. 5A; *t*_(10)_*=* 2.638, *P* = 0.0372, Unpaired Welch’s *t-*test). In the subiliac lymph node, in accordance with the unaltered expression level of MHC class II gene *H2-Aa* (Fig. 4M), *Cd40lg* expression was unchanged (Fig. 5B; *t*_(10)_*=* 0.3956, *P* = 0.7007, Unpaired Student’s *t-*test). Next, the gene expression levels of the Th cell subset markers *Tbet* (Th1), *Gata3* (Th2), *Rorc* (Th17), and *Foxp3* (Treg) were analyzed in the spinal cord. The expression level of the Th17 marker showed a strong, but non-significant, decreasing trend (Fig. 5C; *t*_(12)_*=* 0.9191, *P* = 0.3761, Unpaired Student’s *t-*test for *Tbet*; *t*_(12)_*=* 1.696, *P* = 0.1156, Unpaired Student’s *t-*test for *Gata3*; *t*_(12)_*=* 1.841, *P* = 0.0904, Unpaired Student’s *t-*test for *Rorc*; *t*_(12)_*=* 1.891, *P* = 0.083, Unpaired Student’s *t-*test for *Foxp3*). In the subiliac lymph node and spleen, none of the subset markers were changed, indicating comparable priming of CD4^+^ T cells in the periphery (Fig. 5D; *t*_(10)_*=* 1.730, *P* = 0.1143, Unpaired Student’s *t-*test for *Tbet*; *t*_(10)_*=* 1.210, *P* = 0.2542, Unpaired Student’s *t-*test for *Gata3*; *t*_(10)_*=* 1.200, *P* = 0.2578, Unpaired Student’s *t-*test for *Rorc*; *t*_(10)_*=* 1.169, *P* = 0.2696, Unpaired Student’s *t-*test for *Foxp3*; Fig. 5E; *t*_(10)_*=* 0.006770, *P* = 0.9948, Unpaired Welch’s *t-*test for *Tbet*; *t*_(10)_*=* 1.271, *P* = 0.2536, Unpaired Welch’s *t-*test for *Gata3*; *t*_(10)_*=* 1.040, *P* = 0.3228, Unpaired Student’s *t-*test for *Rorc*; *t*_(10)_*=* 0.1149, *P* = 0.9108, Unpaired Student’s *t-*test for *Foxp3*). Therefore, we focused on IL17-producing cells and further investigation was performed using immunohistochemistry. The number of CD3^+^ T cells and CD3^+^IL17^+^ Th17 cells was consistently reduced in the spinal cords of the OPC-depleted group (Fig. 5F-H; *t*_(16)_*=* 2.206, *P* = 0.0424, Unpaired Student’s *t-*test for Fig. 5G; *t*_(16)_*=* 2.212, *P* = 0.0418, Unpaired Student’s *t-*test for Fig. 5H); however, the percentage of CD3^+^IL17^+^ cells (as a proportion of total CD3^+^ cells) was not altered between the groups (Fig. 5I; *t*_(16)_*=* 0.4628, *P* = 0.6498, Unpaired Student’s *t-*test).

**Figure 5.**
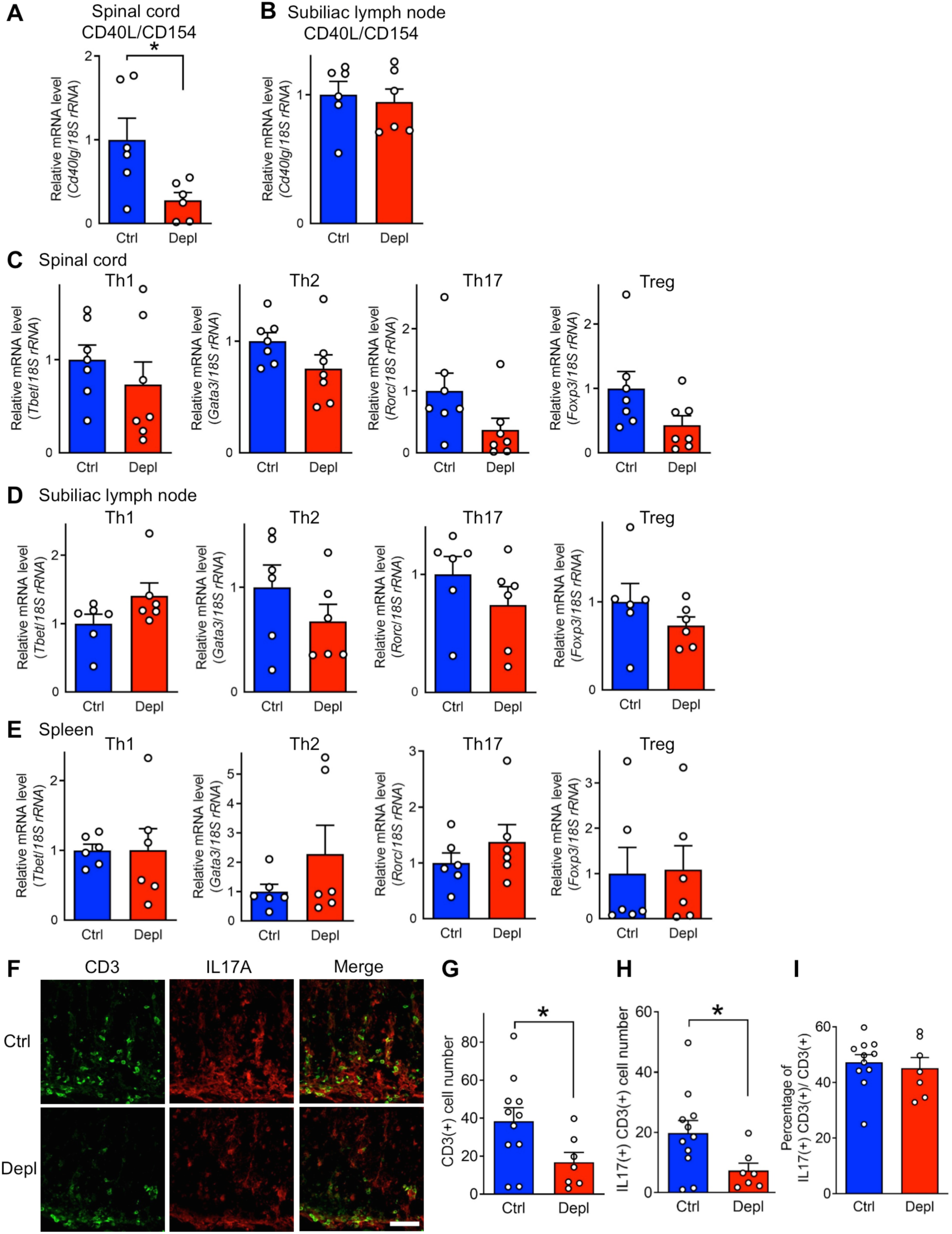
Th17 cell number is decreased in the spinal cord after OPC depletion in the acute phase. ***A, B***. Quantitative PCR analysis of *Cd40lg (CD154)* in the spinal cord (***A***) and subiliac lymph node (***B***) 4 days after the onset of EAE. *n* = 6 mice. ***C***. Quantitative PCR analysis of Th subset markers *Tbet* (Th1), *Gata3* (Th2), *Rorc* (Th17), and *Foxp3* (Treg) in the spinal cord at 4 days after the onset of EAE. *n* = 7 mice. ***D, E***. Quantitative PCR analysis of Th subset markers *Tbet* (Th1), *Gata3* (Th2), *Rorc* (Th17), and *Foxp3* (Treg) in the subiliac lymph node (***D***) and spleen (***E***) 4 days after EAE onset. *n* = 6 mice. ***F-I***. Coimmunostaining of CD3 and IL17A in the spinal cord four days after EAE onset. Representative images are presented in (***F***). The numbers of CD3^+^ cells (***G***) and CD3^+^IL17^+^ cells (***H***), and the percentage of CD3^+^IL17^+^ cells (as a proportion of total CD3^+^) cells (***I***) were analyzed. Scale bar, 50 µm. *n* = 7–11 mice. \**P* < 0.05. Data are expressed as mean ± SEM.

### 3.4 Reduced activation of macrophage was associated with ameliorated EAE in OPC-depleted mice

The decrease in T-cell number in the spinal cord samples from the OPC-depleted group (Fig. 5) may be due to reduced MHC class II expression (Fig. 4K); therefore, the cell type responsible for the change was assessed. MHC class II was mainly expressed by Iba1^+^ cells in spinal cord lesions (Fig. 6A). Although the relative intensity of Iba1 was not significantly different between the groups (Fig. 6B; *t*_(16)_*=* 1.477, *P* = 0.1591, Unpaired Student’s *t-*test), the relative intensity of MHC class II decreased in the spinal cord white matter of the OPC-depleted group (Fig. 6C; *t*_(16)_*=* 2.312, *P* = 0.0344, Unpaired Student’s *t-*test). Previous studies have reported MHC class II expression in OPCs (Falcão et al., 2018; Zveik et al., 2022); however, we did not detect PDGFRα^+^MHCII^+^ cells in WT EAE mice (Fig. 6D). Rather, some PDGFRα^+^ cells were localized adjacent to MHC class II^+^ cells, suggesting a possible interaction between them (Fig. 6D). To further examine the characteristics of Iba1^+^MHC class II^+^ cells, cell type marker expression was investigated in WT EAE mice. Co-expression of MHC class II in Iba1^+^ cells was confirmed in the acute white matter lesions of WT mice (Fig. 6E). Iba1^+^MHC class II^+^ cells in the lesions expressed high levels of CD45, whereas most Iba1^+^ cells in the rim area were MHC class II^−^ and expressed low levels of CD45 (Fig. 6E). As microglia are CD45^low^ and macrophages are CD45^high^, the Iba1^+^MHC class II^+^ cells in the lesions are seemingly macrophages. The Iba1^+^MHC class II^+^ cells also expressed CD11c (Fig. 6F), which is consistent with a previous study showing that monocyte-derived antigen-presenting cells co-express *Cd11c* (Monaghan et al., 2019). These results suggest that MHC class II expression in macrophages is decreased in the spinal cords of OPC-depleted mice, leading to a reduced expansion of T cells and Th17 cells.

**Figure 6.**
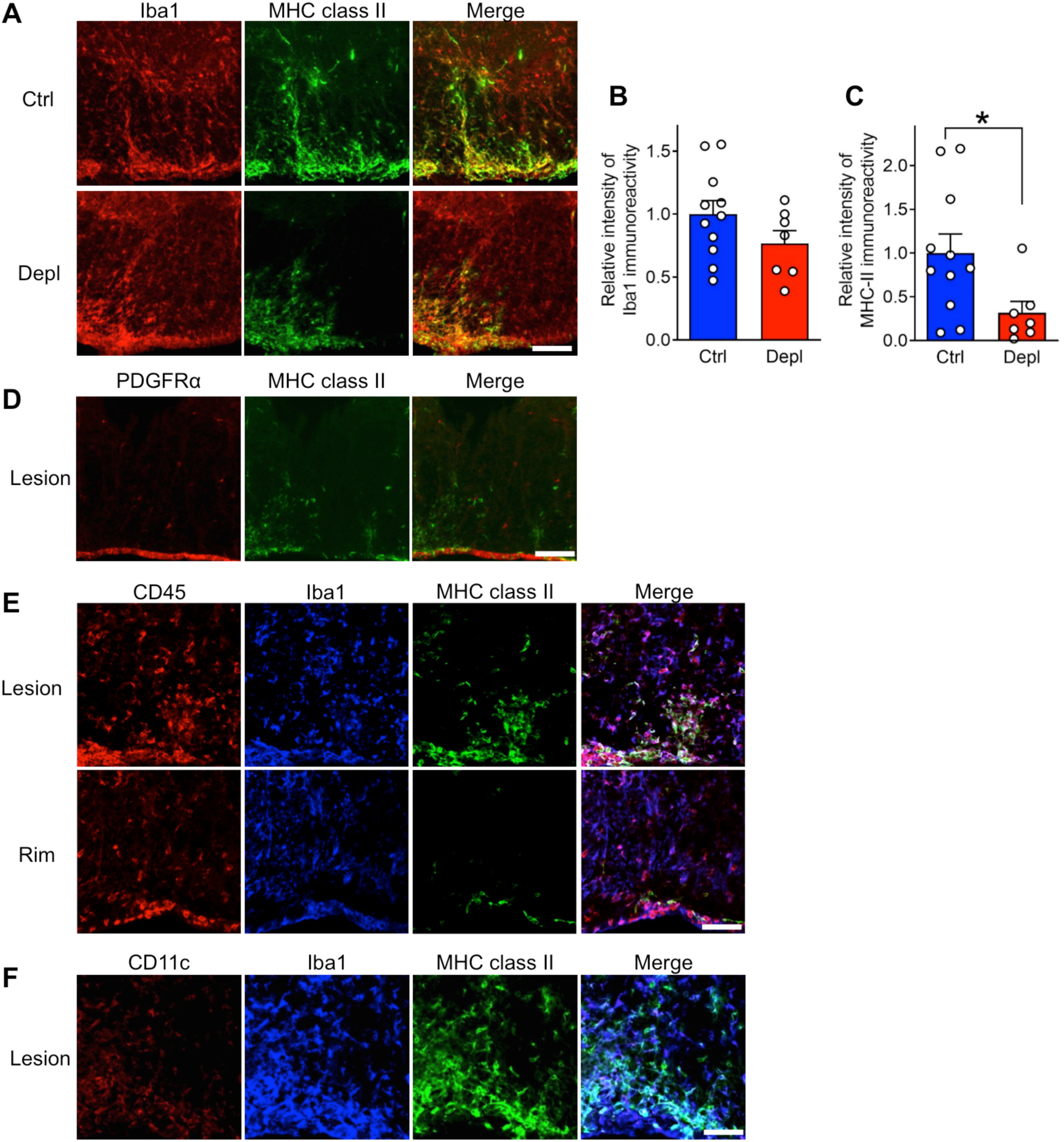
MHC class II expression in macrophages is decreased in the spinal cord after OPC depletion in acute phase. ***A-C***. Representative images of co-immunostaining of Iba1 and MHC class II in the spinal cord at 4 days after the onset of EAE (***A***) and quantification of relative fluorescent intensity of Iba1 (***B***) and MHC class II (***C***) in the white matter. Scale bar, 100 µm. *n* = 7–11 mice. ***D***. Co-immunostaining of PDGFRα and MHC class II in the acute phase (2 days after onset) white matter lesion in the spinal cord of wildtype mice. Scale bar, 100 µm. ***E***. Co-immunostaining of CD45, Iba1, and MHC class II in the acute phase (2 days after onset) white matter lesion and rim in the spinal cord of wildtype mice. Scale bar, 50 µm. ***F***. Co-immunostaining of CD11c, Iba1, and MHC class II in the acute phase (2 days after onset) white matter lesion in the spinal cord of wildtype mice. Scale bar, 50 µm. \**P* < 0.05. Data are expressed as mean ± SEM.

## 4 DISCUSSION

In the current study, we found that depletion of PDGFRα^+^ OPCs during the acute phase of EAE improved the clinical score and reduced T cells in the spinal cord, implying the detrimental role of OPCs. Although OPCs are a source of mature oligodendrocytes that contribute to myelin repair, contrary to our initial expectations, OPC depletion in the acute phase did not induce demyelination but rather attenuated demyelination. T cells in the spinal cords of OPC-depleted mice were decreased, and this correlated with reduced MHC class II expression on macrophages. Although future research is needed to determine the precise mechanisms, this study indicates that in the spinal cord during EAE, OPCs may play a role in enhancing activation of antigen-presenting macrophages, which may lead to T-cell reactivation.

In this study, we focused on the role of undifferentiated OPCs during the short time window following EAE symptom onset, rather than the long-term role of OPCs. Rivers et al. (2008) demonstrated that it takes more than two months for 70 % of PDGFRα^+^ OPCs to differentiate into myelinating oligodendrocytes in P45 healthy adult mice. As the experimental period in the current study was comparatively short, selective depletion of immature OPCs without loss of mature oligodendrocytes or myelin was achieved. It is possible that if the duration of OPC depletion was longer or tamoxifen was administered at earlier timepoint, the results could have been the opposite of those observed here: mature oligodendrocytes reduced and myelin repair inhibited.

The results of this study, in which OPCs regulate immune function in a pathology-progressive manner in the early stages of EAE, raise questions about the assumption that OPCs are a protective cell type in EAE. OPCs effectively differentiate into oligodendrocytes and contribute to remyelination in lysophosphatidylcholine- or cuprizone-induced demyelination (Baxi et al., 2017; Serwanski et al., 2018). On the other hand, in EAE, where demyelination is caused by strong inflammation, the percentage of OPCs differentiate is low presumably because of the inhibitory environments for remyelination (Coppolino et al., 2018). The present study indicates that in acute EAE, OPCs contribute more to neuroinflammation than to myelin repair. As OPCs may be heterogenous population, differentiation and immune modulation may be mediated by different subsets. Recently, it was reported that immune-related genes including MHC molecules are upregulated in a subset of OPCs sorted from MS or EAE brain (Falcão et al., 2018; Jäkel et al., 2019; Kirby et al., 2019; Meijer M, et al., 2022). The current result is different in that MHC class II was not expressed by OPCs and it seems that OPCs themselves do not present antigens but instead promote the function of antigen presenting cells. However, this is the first in vivo demonstration of immunomodulatory role of OPCs in EAE, highlighting the need for further research on potential interaction of OPCs and immune cells. As we cannot rule out the possibility that OPC depletion itself triggers inflammation and affects the function of other cells, next study should target specific molecules on OPCs.

The day of onset did not vary between the groups of transgenic mice (Fig. 1). On the other hand, the day of onset was delayed by about 1 week compared to previous study (Tsutsui et al., 2018). What caused the delay is unclear, but it is reasonable to infer that administration of tamoxifen on 10-12 days post immunization delayed the day of onset given that tamoxifen suppresses EAE symptoms (Bebo et al., 2009).

Why did OPC depletion ameliorate demyelination during acute EAE? Our results show that MHC class II, which was mostly expressed in macrophages, was decreased in the spinal cord samples from OPC-depleted EAE mice (Fig. 4K, 6A, 6C). Macrophages are derived from monocytes and not only directly phagocytose myelin and initiate demyelination (Yamasaki et al., 2014) but also present antigens and produce inflammatory cytokines (Wang et al., 2019). Antigen presentation via MHC class II is essential for reactivating T cells and promoting their expansion in the CNS (Chastain et al., 2011; Dong and Yong, 2019). Recently, experiments using *Cx3cr1^CreERT2^* and *Cx3cr1^CreER^*mice showed that MHC class II-dependent antigen presentation in the CNS is mediated by bone-marrow-derived cells and not by embryonically derived microglia and border-associated macrophages (Wolf et al., 2018, Mundt et al., 2019). Bone-marrow-derived macrophages and dendritic cells could not be distinguished in this system (Wolf et al., 2018) and both may act as antigen-presenting cells. In the present study, Iba1^+^MHC class II^+^ macrophages co-expressed CD11c (Fig. 6F); therefore, they could also have properties of dendritic cells. Taken together, MHC class II expression on macrophages may contribute to reactivation of T cells (Fig. 5), but as macrophages and T cells interact bi-directionally, further mechanistic studies are needed.

To date, little is known regarding the modulation of macrophage activation by OPCs. The current results imply that OPCs in acute EAE facilitated macrophage activation and increased MHC class II levels in the spinal cord. The transcription factor NFAT5 positively regulates the transactivator CIITA and downstream MHC class II in macrophages (Buxadé et al., 2018). NFAT5 expression is enhanced by IFNγ and Toll-like receptor (TLR) stimulation (Lee et al., 2019). Therefore, OPCs may be involved in the activation of IFNγ receptor or TLR signalings in macrophages. We observed OPCs in proximity to MHC class II-expressing cells in the acute lesions (Fig. 6D); therefore, direct cell–cell contact with OPCs within the spinal cord parenchyma may promote these signaling pathways in macrophages. The interaction between OPCs and macrophages has been postulated in a study demonstrating that TLR4 on macrophages could bind to NG2 proteoglycan on OPCs (Hayakawa et al., 2016). Another possibility is that cytokines produced by OPCs promote macrophage activation. OPCs express various cytokines in pathologies, including IL1β (Moyon et al., 2015; Zeis et al., 2016), which enhances NFAT5 expression in macrophages (Choi et al., 2017). Future research should focus on the mechanisms by which OPCs facilitate MHC class II expression by macrophages.

This study suggests that suppressing macrophage activation in the CNS can attenuate neuroinflammation by inhibiting subsequent T-cell expansion. The pathological role of macrophages was also shown in a previous report where monocyte depletion after the onset of EAE symptoms diminished the peak clinical score and reduced CD4^+^ T cells in the CNS (Moreno et al., 2016). Generally, current MS therapies target T and B cells, although some reports suggested that these drugs also attenuate macrophage activation (Mishra et al., 2016; Nally et al., 2019), which could be important for efficacy of the treatments. Further studies of the molecular mechanisms by which OPCs activate macrophages may reveal new therapeutic strategies for MS.

Some OPCs express matrix metallopeptidase 9 (Seo et al., 2013) and are involved in the disruption of the BBB in pathogenic conditions (Maki, 2017). Dysfunctionally clustered perivascular OPCs evict astrocyte endfeet from BBB, disrupt tight junctions between endothelial cells, and increase BBB permeability (Niu et al., 2019). Therefore, because OPCs might be involved with BBB/BSCB integrity, we investigated the expression levels of BSCB tight junction proteins and adhesion molecules; however, no changes were observed (Fig. 4C–H). Thus, the reduced numbers of T cells and Th17 cells in the spinal cords of OPC-depleted mice may not be due to changes in BSCB integrity.

We used PDGFRα as a marker for OPCs, but PDGFRα is also expressed by CNS fibroblasts during the chronic phase of EAE (Dorrier et al., 2021). Consistent with previous studies (Yahn et al., 2020; Dorrier et al., 2021), only small numbers of PDGFRα^+^collagen I^+^ fibroblasts were present in acute lesions, whereas fibroblast number was increased in chronic lesions (Fig. 3). Considering previous reports that depletion of collagen-positive fibroblasts augmented the EAE clinical score only in the chronic stage and there was no difference in the number of T cells in the spinal cord (Dorrier et al., 2021), the present results are different from that of fibroblast depletion. Another OPC marker, NG2, is expressed in activated pericytes (Girolamo et al., 2019) and macrophages (Moransard et al., 2011) in EAE and lacks specificity. Therefore, more specific OPC removal strategies, such as Sox10^+^PDGFRα^+^ OPCs depletion (Xing et al., 2023) and Olig2^+^ PDGFRα^+^OPCs depletion (Brousse et al., 2023), are needed to better understand the role of OPCs under EAE.

In summary, this study demonstrated that OPC deletion during the acute phase of EAE reduces the paralysis and demyelination via suppression of macrophage activation and T-cell reactivation in the CNS. The full molecular mechanism remains unelucidated; however, the current results provide new insight that OPCs function not only as the source for myelinating oligodendrocytes but they also play roles in immune regulation in the context of acute neuroinflammation.

## ACKNOWLEDGMENTS

This work was supported by Grants-in-Aid for Scientific Research (KAKENHI) from MEXT/JSPS (to H.S., JP19K22494, 23H02639), Grants-in-Aid for JSPS Fellows (to K.O., JP19J23074), and also by the Takeda Science Foundation and the Uehara memorial Foundation (to H.S.)

## DATA AVAILABILITY STATEMENT

The data that support the findings of this study are available from the corresponding author upon reasonable request.

## REFERENCES

Barres, BA., Hart, IK., Coles, HS., Burne, JF., Voyvodic, JT., Richardson, & WD., Raff. MC. (1992). Cell death and control of cell survival in the oligodendrocyte lineage. Cell, 70, 31–46. 10.1016/0092-8674(92)90531-g

Baxi, EG., DeBruin, J., Jin, J., Strasburger, HJ., Smith, MD., … & Calabresi, PA. (2017). Lineage tracing reveals dynamic changes in oligodendrocyte precursor cells following cuprizone-induced demyelination. Glia, 65, 2087–2098. 10.1002/glia.23229

Bebo, BF Jr., Dehghani, B., Foster, S., Kurniawan, A., Lopez, FJ., & Sherman, LS. (2009). Treatment with selective estrogen receptor modulators regulates myelin specific T-cells and suppresses experimental autoimmune encephalomyelitis. Glia, 57, 777–90. 10.1002/glia.20805

Brousse, B., Mercier, O., Magalon, K., Gubellini, P., Malapert, P., Cayre, M., & Durbec, P. (2023). Characterization of a new mouse line triggering transient oligodendrocyte progenitor depletion. Scientific reports, 13(1), 21959. 10.1038/s41598-023-48926-4

Buxadé, M., Huerga Encabo, H., Riera-Borrull, M., Quintana-Gallardo, L., López-Cotarelo, P., Tellechea, M., … & López-Rodríguez, C. (2018). Macrophage-specific MHCII expression is regulated by a remote Ciita enhancer controlled by NFAT5. J Exp Med, 215, 2901–2918. 10.1084/jem.20180314

Chastain, EM., Duncan, DS., Rodgers, JM., & Miller, SD. (2011). The role of antigen presenting cells in multiple sclerosis. Biochim Biophys Acta, 1812, 265–274. 10.1016/j.bbadis.2010.07.008

Choi, S., You, S., Kim, D., Choi, SY., Kwon, HM., Kim, HS., & Kim, WU. (2017). Transcription factor NFAT5 promotes macrophage survival in rheumatoid arthritis. J Clin Invest, 127, 954–969. 10.1172/JCI87880

Coppolino, GT., Marangon, D., Negri, C., Menichetti, G., Fumagalli, M., Gelosa, P., … & Abbracchio, MP. (2018). Differential local tissue permissiveness influences the final fate of GPR17-expressing oligodendrocyte precursors in two distinct models of demyelination. Glia, 66, 1118–1130. 10.1002/glia.23305

Dong, Y., & Yong, VW. (2019). When encephalitogenic T cells collaborate with microglia in multiple sclerosis. Nat Rev Neurol, 15, 704–717. 10.1038/s41582-019-0253-6

Dorrier, CE., Aran, D., Haenelt, EA., Sheehy, RN., Hoi, KK., Pintarić, L., … & Daneman, R. (2021). CNS fibroblasts form a fibrotic scar in response to immune cell infiltration. Nat Neurosci, 24, 234–244. 10.1038/s41593-020-00770-9

Falcão, AM., van Bruggen, D., Marques, S., Meijer, M., Jäkel, S., Agirre, E., … & Castelo-Branco, G. (2018). Disease-specific oligodendrocyte lineage cells arise in multiple sclerosis. Nat Med, 24, 1837–1844. 10.1038/s41591-018-0236-y

Franklin, RJM., & Ffrench-Constant, C. (2017). Regenerating CNS myelin - from mechanisms to experimental medicines. Nat Rev Neurosci, 18, 753–769. 10.1038/nrn.2017.136

Fünfschilling, U., Supplie, LM., Mahad, D., Boretius, S., Saab, AS., Edgar, J., … & Nave, KA. (2012). Glycolytic oligodendrocytes maintain myelin and long-term axonal integrity. Nature, 485, 517–521. 10.1038/nature11007

Girolamo, F., Errede, M., Longo, G., Annese, T., Alias, C., Ferrara, G., … & Virgintino, D. (2019). Defining the role of NG2-expressing cells in experimental models of multiple sclerosis. A biofunctional analysis of the neurovascular unit in wild type and NG2 null mice. PLoS One, 14, e0213508. 10.1371/journal.pone.0213508

Hayakawa, K., Pham, LD., Seo, JH., Miyamoto, N., Maki, T., Terasaki, Y., … & Lo, EH. (2016). CD200 restrains macrophage attack on oligodendrocyte precursors via toll-like receptor 4 downregulation. J Cereb Blood Flow Metab, 36, 781–793. 10.1177/0271678X15606148

Jäkel, S., Agirre, E., Mendanha Falcão, A., van Bruggen, D., Lee, KW., Knuesel, I., … & Castelo-Branco, G. (2019). Altered human oligodendrocyte heterogeneity in multiple sclerosis. Nature, 566, 543–547. 10.1038/s41586-019-0903-2

Kawakami, N., Lassmann, S., Li, Z., Odoardi, F., Ritter, T., Ziemssen, T., … & Flügel, A. (2004). The activation status of neuroantigen-specific T cells in the target organ determines the clinical outcome of autoimmune encephalomyelitis. J Exp Med, 199, 185–197. 10.1084/jem.20031064

Kirby, L., Jin, J., Cardona, JG., Smith, MD., Martin, KA., Wang, J., … & Calabresi, PA. (2019). Oligodendrocyte precursor cells present antigen and are cytotoxic targets in inflammatory demyelination. Nat Commun, 10, 3887. 10.1038/s41467-019-11638-3

Lee, N., Kim, D., & Kim, WU. (2019). Role of NFAT5 in the Immune System and Pathogenesis of Autoimmune Diseases. Front Immunol, 10, 270. 10.3389/fimmu.2019.00270

Liu, X., Xin, D. E., Zhong, X., Zhao, C., Li, Z., Zhang, L., … & Lu, Q. R. (2024). Small-molecule-induced epigenetic rejuvenation promotes SREBP condensation and overcomes barriers to CNS myelin regeneration. Cell, 187(10), 2465–2484.e22. 10.1016/j.cell.2024.04.005

Maki, T. (2017). Novel roles of oligodendrocyte precursor cells in the developing and damaged brain. Clin Exp Neuroimmunol, 8, 33–42. 10.1111/cen3.12358

Meijer, M., Agirre, E., Kabbe, M., van Tuijn, C. A., Heskol, A., Zheng, C., … & Castelo-Branco, G. (2022). Epigenomic priming of immune genes implicates oligodendroglia in multiple sclerosis susceptibility. Neuron, 110(7), 1193–1210.e1113. 10.1016/j.neuron.2021.12.034

Mishra, MK., & Yong, VW. (2016). Myeloid cells - targets of medication in multiple sclerosis. Nat Rev Neurol, 12, 539–551. 10.1038/nrneurol.2016.110

Miyamura, S., Matsuo, N., Nagayasu, K., Shirakawa, H., & Kaneko, S. (2019). Myelin Oligodendrocyte Glycoprotein 35-55 (MOG 35-55)-induced Experimental Autoimmune Encephalomyelitis: A Model of Chronic Multiple Sclerosis. Bio Protoc, 9, e3453. 10.21769/BioProtoc.3453

Monaghan, KL., Zheng, W., Hu, G., & Wan, ECK. (2019). Monocytes and Monocyte-Derived Antigen-Presenting Cells Have Distinct Gene Signatures in Experimental Model of Multiple Sclerosis. Front Immunol, 10, 2779. 10.3389/fimmu.2019.02779

Moransard, M., Dann, A., Staszewski, O., Fontana, A., Prinz, M., & Suter, T. (2011). NG2 expressed by macrophages and oligodendrocyte precursor cells is dispensable in experimental autoimmune encephalomyelitis. Brain, 134, 1315–1330. 10.1093/brain/awr070

Moreno, MA., Burns, T., Yao, P., Miers, L., Pleasure, D., & Soulika, AM. (2016). Therapeutic depletion of monocyte-derived cells protects from long-term axonal loss in experimental autoimmune encephalomyelitis. J Neuroimmunol, 290, 36–46. 10.1016/j.jneuroim.2015.11.004

Moyon, S., Dubessy, AL., Aigrot, MS., Trotter, M., Huang, JK., Dauphinot, L., … & Lubetzki, C. (2015). Demyelination causes adult CNS progenitors to revert to an immature state and express immune cues that support their migration. J Neurosci, 35, 4–20. 10.1523/JNEUROSCI.0849-14.2015

Mundt, S., Mrdjen, D., Utz, SG., Greter, M., Schreiner, B., & Becher, B. (2019). Conventional DCs sample and present myelin antigens in the healthy CNS and allow parenchymal T cell entry to initiate neuroinflammation. Sci Immunol, 4, eaau8380. 10.1126/sciimmunol.aau8380

Nally, FK., De Santi, C., & McCoy, CE. (2019). Nanomodulation of Macrophages in Multiple Sclerosis. Cells, 8, 543. 10.3390/cells8060543

Nicaise, AM., Wagstaff, LJ., Willis, CM., Paisie, C., Chandok, H., Robson, P., … & Crocker, SJ. (2019). Cellular senescence in progenitor cells contributes to diminished remyelination potential in progressive multiple sclerosis. Proc Natl Acad Sci U S A, 116, 9030–9039. 10.1073/pnas.1818348116

Niu, J., Tsai, HH., Hoi, KK., Huang, N., Yu, G., Kim, K., … & Fancy, SPJ. (2019). Aberrant oligodendroglial-vascular interactions disrupt the blood-brain barrier, triggering CNS inflammation. Nat Neurosci, 22, 709–718. 10.1038/s41593-019-0369-4

Planas, R., Metz, I., Martin, R., & Sospedra, M. (2018). Detailed Characterization of T Cell Receptor Repertoires in Multiple Sclerosis Brain Lesions. Front Immunol, 9, 509. 10.3389/fimmu.2018.00509

Rivers, LE., Young, KM., Rizzi, M., Jamen, F., Psachoulia, K., Wade, A., … & Richardson, WD. (2008). PDGFRA/NG2 glia generate myelinating oligodendrocytes and piriform projection neurons in adult mice. Nat Neurosci, 11, 1392–1401. 10.1038/nn.2220

Seo, JH., Miyamoto, N., Hayakawa, K., Pham, LD., Maki, T., Ayata, C., … & Arai, K. (2013). Oligodendrocyte precursors induce early blood-brain barrier opening after white matter injury. J Clin Invest, 123, 782–786. 10.1172/JCI65863

Serwanski, DR., Rasmussen, AL., Brunquell, CB., Perkins, SS., & Nishiyama, A. (2018). Sequential Contribution of Parenchymal and Neural Stem Cell-Derived Oligodendrocyte Precursor Cells toward Remyelination. Neuroglia, 1, 91–105. 10.3390/neuroglia1010008

Tiane, A., Schepers, M., Reijnders, R. A., van Veggel, L., Chenine, S., Rombaut, B., … & Vanmierlo, T. (2023). From methylation to myelination: epigenomic and transcriptomic profiling of chronic inactive demyelinated multiple sclerosis lesions. Acta neuropathologica, 146(2), 283–299. 10.1007/s00401-023-02596-8

Tsutsui, M., Hirase, R., Miyamura, S., Nagayasu, K., Nakagawa, T., Mori, Y., … & Kaneko, S. (2018). TRPM2 Exacerbates Central Nervous System Inflammation in Experimental Autoimmune Encephalomyelitis by Increasing Production of CXCL2 Chemokines. J Neurosci, 38, 8484–8495. 10.1523/JNEUROSCI.2203-17.2018

Wang, J., Wang, J., Wang, J., Yang, B., Weng, Q., & He, Q. (2019). Targeting Microglia and Macrophages: A Potential Treatment Strategy for Multiple Sclerosis. Front Pharmacol, 10, 286. 10.3389/fphar.2019.00286

Wolf, Y., Shemer, A., Levy-Efrati, L., Gross, M., Kim, JS., Engel, A., … & Jung, S. (2018). Microglial MHC class II is dispensable for experimental autoimmune encephalomyelitis and cuprizone-induced demyelination. Eur J Immunol, 48, 1308–1318. 10.1002/eji.201847540

Xing, Y. L., Poh, J., Chuang, B. H. A., Moradi, K., Mitew, S., Richardson, W. … & Merson, T. D. (2023). High-efficiency pharmacogenetic ablation of oligodendrocyte progenitor cells in the adult mouse CNS. Cell reports methods, 3(2), 100414. 10.1016/j.crmeth.2023.100414

Yahn, SL., Li, J., Goo, I., Gao, H., Brambilla, R., & Lee, JK. (2020). Fibrotic scar after experimental autoimmune encephalomyelitis inhibits oligodendrocyte differentiation. Neurobiol Dis, 134, 104674. 10.1016/j.nbd.2019.104674

Yamasaki, R., Lu, H., Butovsky, O., Ohno, N., Rietsch, AM., Cialic, R., … & Ransohoff, RM. (2014). Differential roles of microglia and monocytes in the inflamed central nervous system. J Exp Med, 211, 1533–1549. 10.1084/jem.20132477

Yeung, MSY., Djelloul, M., Steiner, E., Bernard, S., Salehpour, M., Possnert, G., … & Frisén, J. (2019). Dynamics of oligodendrocyte generation in multiple sclerosis. Nature, 566, 538–542. 10.1038/s41586-018-0842-3

Zeis, T., Enz, L., & Schaeren-Wiemers, N. (2016). The immunomodulatory oligodendrocyte. Brain Res, 1641, 139–148. 10.1016/j.brainres.2015.09.021

Zveik, O., Fainstein, N., Rechtman, A., Haham, N., Ganz, T., Lavon, I., … & Vaknin-Dembinsky, A. (2022). Cerebrospinal fluid of progressive multiple sclerosis patients reduces differentiation and immune functions of oligodendrocyte progenitor cells. Glia, 70, 1191–1209. 10.1002/glia.24165

